# *Candida albicans* genetic background influences mean and heterogeneity of drug responses and genome stability during evolution to fluconazole

**DOI:** 10.1101/360347

**Authors:** Aleeza C. Gerstein, Judith Berman

**Author notes:** Correspondence: Aleeza C. Gerstein, Judith Berman.

## Abstract

The importance of within-species diversity in determining the evolutionary potential of a population to evolve drug resistance or tolerance is not well understood, including in eukaryotic pathogens. To examine the influence of genetic background, we evolved replicates of twenty different clinical isolates of *Candida albicans,* a human fungal pathogen, in fluconazole, the commonly used antifungal drug. The isolates hailed from the major *C. albicans* clades and had different initial levels of drug resistance and tolerance to the drug. The majority of replicates rapidly increased in fitness in the evolutionary environment, with the degree of improvement inversely correlated with ancestral strain fitness in the drug. Improvement was largely restricted to up to the evolutionary level of drug: only 4% of the evolved replicates increased resistance (MIC) above the evolutionary level of drug. Prevalent changes were altered levels of drug tolerance (slow growth of a subpopulation of cells at drug concentrations above the MIC) and increased diversity of genome size. The prevalence and predominant direction of these changes differed in a strain-specific manner but neither correlated directly with ancestral fitness or improvement in fitness. Rather, low ancestral strain fitness was correlated with high levels of heterogeneity in fitness, tolerance, and genome size among evolved replicates. Thus, ancestral strain background is an important determinant in mean improvement to the evolutionary environment as well as the diversity of evolved phenotypes, and the range of possible responses of a pathogen to an antimicrobial drug cannot be captured by in-depth study of a single strain background.

**Importance:** Antimicrobial resistance is an evolutionary phenomenon with clinical implications. We tested how replicates from diverse strains of *Candida albicans*, a prevalent human fungal pathogen, evolve in the commonly-prescribed antifungal drug fluconazole. Replicates on average increased in fitness in the level of drug they were evolved to, with the least fit ancestral strains improving the most. Very few replicates increased resistance above the drug level they were evolved in. Notably, many replicates increased in genome size and changed in drug tolerance (a drug response where a subpopulation of cells grow slowly in high levels of drug) and variability among replicates in fitness, tolerance and genome size was higher in strains that initially were more sensitive to the drug. Genetic background influenced the average degree of adaptation and the evolved variability of many phenotypes, highlighting that different strains from the same species may respond and adapt very differently during adaptation.

## Introduction

In eukaryotic microbes, the responses to severe stresses, including exposure to antimicrobial drugs, can occur through genetic changes that arise within susceptible microbial populations and spread via conventional evolutionary processes or via physiological responses that modulate the ability of cells to survive and grow in the presence of the stress. Drug resistance, measured as reduced susceptibility, assessed as a higher minimal inhibitory concentration (MIC). For fungal pathogens, broth microdilution assays, e-test strips or disk diffusion assays are assessed after 24h of growth in a set range of drug concentrations (1). Antifungal drug tolerance, a property distinct from drug resistance (2, 3), is the ability of some cells in a strain to grow slowly in the presence of a drug at concentrations above the MIC. In tolerant strains, the subpopulation of cells that grow (generally from 10-90% of cells, depending on the strain) is usually evident when growth is assessed after 48 h or longer in the drug (reviewed in 4). Hence, tolerant strains have susceptible MIC levels (5, 6) and have often been termed ‘resistant’ in assays that are allowed to grow for longer periods of time and/or in spot assays that measure partial growth. Clinically, the failure to clear infections is more likely when the infecting strain is drug-resistant. High levels of drug tolerance also may influence infection clearance (3, 7), although more studies that use quantitative criteria to definitively measure tolerance levels and distinguish tolerance from resistant isolates are needed (reviewed in 4).

Factors that influence the change in antifungal resistance and tolerance levels in pathogenic fungi have not been well elucidated. Unlike prokaryotes, which often acquire new traits horizontally via plasmids, eukaryotic pathogens primarily acquire new traits vertically via *de novo* mutations, chromosome-scale changes in copy number (ploidy) or allele frequency, and recombination events. Changes in either genome-wide ploidy (the number of sets of homologous chromosome) or aneuploidy (the gain or loss of individual chromosomes) arise in laboratory populations subjected to stress (8–10) or passaged through mice (11, 12). Aneuploidy is also found in some clinical isolates of *C. albicans, Candida glabrata* and *Saccharomyces cerevisiae* (13, 14) as well as in wild type environmental isolates of *S. cerevisiae* (15, 16).

Changes in ploidy arise more frequently than point mutations (17). In addition, and are especially prevalent in strains exposed to azole drug stress (18–22). Indeed, exposure to fluconazole promotes karyotypic change by inducing unconventional cell cycle events in a subpopulation of cells (23, 24). Hence, fluconazole can both drive and select for karyotypic variation within and among populations. Furthermore, the frequency of aneuploidy among clinical isolates may be underestimated, as strain isolation methods (i.e., multiple growth cycles in rich medium) may impose a fitness cost that selects against aneuploid isolates. A mechanistic link between drug resistance and specific aneuploidies exists in multiple pathogenic fungal species (e.g., *C. albicans,* 22, 25–28; *Cryptococcus neoformans*, 29, 30; suggested in *Candida auris,* 31, 32). Because fluconazole tolerance and drug resistance are distinct and tolerance is sensitive to inhibitors that do not affect resistance (2, 3), it follows that antifungal tolerance likely evolves via a different subset of genes than resistance, although it is also possible that there is overlap between genes involved in tolerance and resistance.

The importance of genetic background in the link between genotype and phenotype for *de novo* acquired mutations is becoming appreciated in both laboratory and natural settings (33–35), and references within). For example, the phenotype of deleted or repressed genes can vary significantly in different backgrounds (36–38). Moreover, closely related strains can differ in the classification of genes that are essential for viability under a specific growth condition (39). The same is true of mutations classified as beneficial in one strain background (including different mutations in the same gene, 40), which can be neutral or deleterious in other backgrounds (e.g., 41–44). Different strain backgrounds also may exhibit different mutation rates, thereby affecting the frequency with which genetic variation arises (45, 46). Thus, the genetic background of a population likely influences the mutations available for adaptation, and hence the trajectory of evolution. Furthermore, the constraints on variation in the degree of intra- and inter-population heterogeneity are likely to differ in different genetic backgrounds.

Here we explored the interplay between genetic background, fitness, drug resistance and drug tolerance, and karyotypic variation by following the evolutionary trajectories of replicates from 20 diverse *C. albicans* strains for 100 generations of evolution in 1µg/ml of fluconazole. We found that the majority of replicates acquired the ability to grow more rapidly in the evolutionary level of drug, with the degree of improvement inversely correlated with ancestral strain fitness. While very few replicates from any background acquired clinical levels of drug resistance, changes in tolerance and ploidy were prevalent, especially in strains with low ancestral fitness. We find that drug tolerance is an evolvable phenotype and that changed in the majority of strains. Importantly, evolved variation in fitness, tolerance, and genome size among replicate evolved lines is inversely correlated with ancestral strain fitness. Thus initial strain fitness provides a link between strain genetic background and the acquisition of phenotypic and genotypic diversity among replicate populations adapting to fluconazole.

## Results

We evolved replicate lines from 20 clinical strains of *C. albicans* that span the phylogenetic diversity of the species (47), and vary in mating type zygosity, geographic origin and site of isolation (Table 1). Twelve replicates of each of the 20 strains were evolved in parallel in fluconazole (YPD + 1 μg/mL FLC) for 100 generations (20 strains x 12 replicates per strain = 240 replicates), by serial passaging at 1:1000 dilution every 72 hours for 10 passages. Optical density (OD) in the evolutionary level of fluconazole was measured for ancestral and evolved populations at 24 and 72 h. Ancestral and evolved fitness was measured as optical density (A_600_) at 24 h and at 72 h. We also measured the drug response phenotypes for ancestral and evolved strains from broth microdilution assays. Resistance to fluconazole was measured as MIC_50_, the concentration of drug at which growth is inhibited by at least 50% relative to growth in a drug-free environment, after 24 h of growth. Tolerance to fluconazole was calculated as the ratio of growth at drug concentrations above the MIC relative to growth without drug after 72 h (the transfer time). Although previous work measured tolerance at 48 h, we found that both timepoints gave very similar results. Ancestral optical density at 24 h and 72 h were correlated (Spearman’s rank correlation, S = 158, p-value < 2.2e-16, rho = 0.88), while ancestral resistance and tolerance were not (Spearman’s rank correlation, S = 1141.8, p-value = 0.55) (Table 1).

**Table 1.**
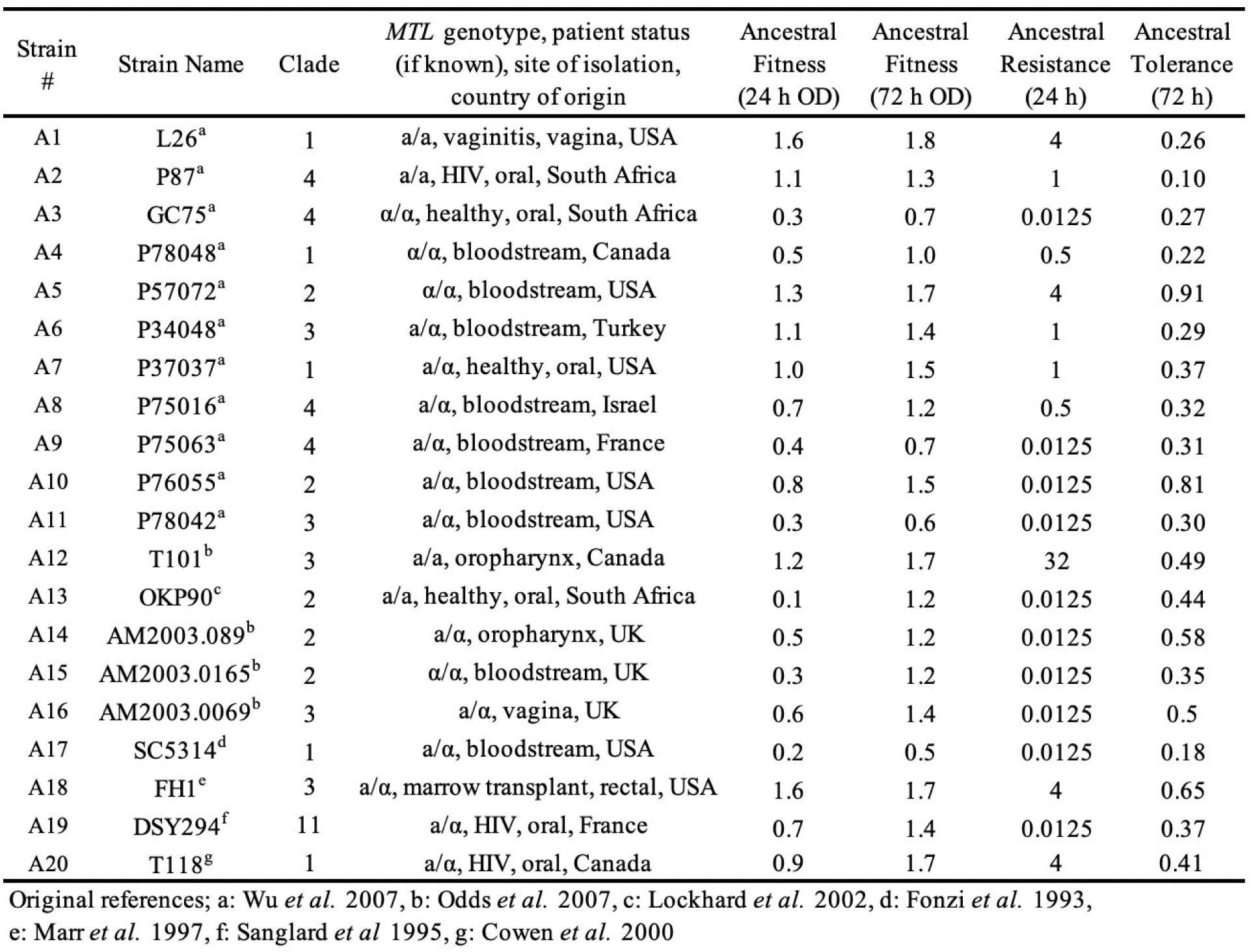
Strains used in this study. Fitness was measured as the optical density (A_600_) in YPD + 1 μg/mL FLC, the evolutionary drug environment. Drug resistance was measured as MIC_50_ at 30 °C by broth microdilution assay. Tolerance was measured as the average optical density at 72 h in the measured drug concentrations of drug above the MIC divided by optical density in the lowest measured drug level.

### Adaptation is influenced by strain background

The majority of replicates evolved significantly higher fitness in the evolutionary drug environment after only 100 generations of adaptation (fitness measured at either 24 and 72 h, Figure 1). The major exception was replicates from A12, the strain with the highest ancestral MIC_50_, which evolved significantly lower 24 h fitness and had no change in 72 h fitness. Four strains with initial MIC ≤ 1 (A2, A9, A10 and A14) also did not acquire improved average fitness at 24 h, though all had significantly increased fitness at 72 h, on average (significance determined as appropriate by parametric or nonparametric t-tests, methods and detailed results in Table S1).

**Figure 1.**
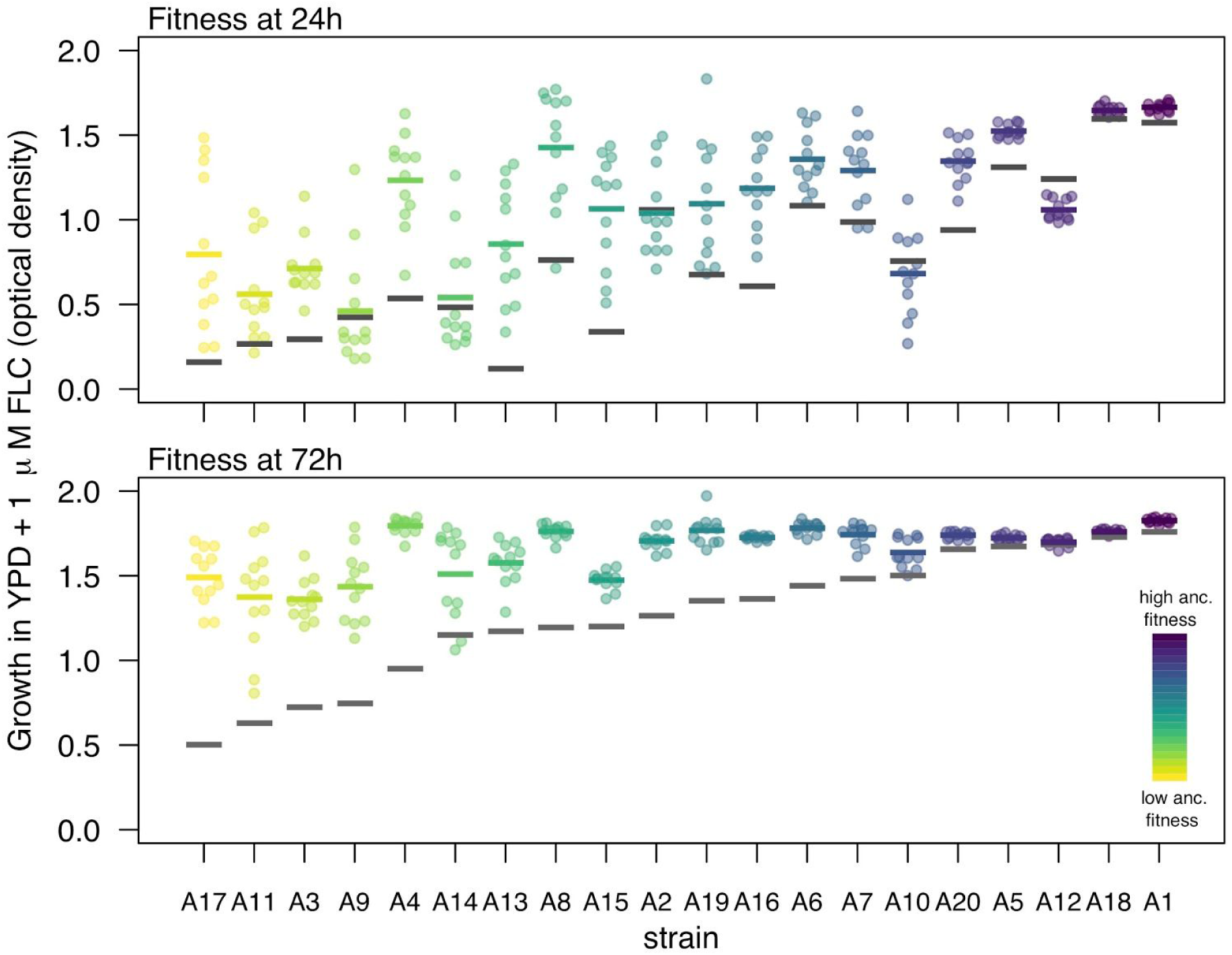
The majority of evolved replicates improved growth in the evolutionary environment after ∼100 generations of evolution. Fitness was measured as growth (optical density) in YPD + 1 μg/mL fluconazole, the evolutionary environment after 24 h (top) and 72 h (bottom). Strains are ordered by ancestral fitness in the evolutionary environment at 72 h. Ancestral fitness (the median growth among 12 ancestral replicates) is indicated for each strain by a grey bar. Each coloured point represents one of twelve evolved replicates while the coloured bars indicate median evolved growth for visual comparison to ancestral growth.

Ancestral strain fitness significantly influenced both the mean fitness improvement after 100 generations and the variability in fitness improvement among the replicates; neither mating type nor clade had a significant effect on these parameters (Figure 2, ANOVA tests; top: 24 h— mean improvement: F_1, 13_ = 7.99, *p* = 0.014, clade: F_4, 13_ = 0.64, *p* = 0.64, *MAT* zygosity: F_1, 13_ = 0.27, *p* = 0.61; variability: F_1, 13_ = 40.94, *p* < 0.0001, clade: F_4, 13_ =0.82, *p* =0.54, *MAT* zygosity: F_1, 13_ = 2.22, *p* = 0.16; 72 h—mean improvement: F_1, 13_ = 158.73, *p* < 0.0001, clade: F_4, 13_ = 1.97, *p* = 0.16, MAT zygosity: F_1, 13_ = 0.25, *p* = 0.63; variability: F_1, 13_ =6.353, *p* < 0.026, clade: F_4, 13_ = 1.01, *p* = 0.44, *MAT* zygosity: F_1, 13_ = 1.13, *p* = 0.31). Additional aspects of strain background that are not accounted for in these models also contributed to evolved fitness, as can be visualized from the deviation of points from the correlation line of fit in Figure 2. The variance among evolved replicates from each strain reflects stochasticity in the evolutionary process. Variance in evolved fitness and the degree of fitness improvement were inversely correlated: strains that were least fit initially and that increased in growth the most on average also had the most variability among replicates from the same strain background (Spearman’s rank correlation test; 24 h: S = 694, *p* = 0.035, rho = 0.48. 72 h: S = 476, *p* = 0.003, rho = 0.64). Overall, strain background influenced both the degree of growth improvement and the variability in improvement among replicates, both mediated in part (but not entirely) by the ancestral growth ability of the strain in the evolutionary environment.

**Figure 2.**
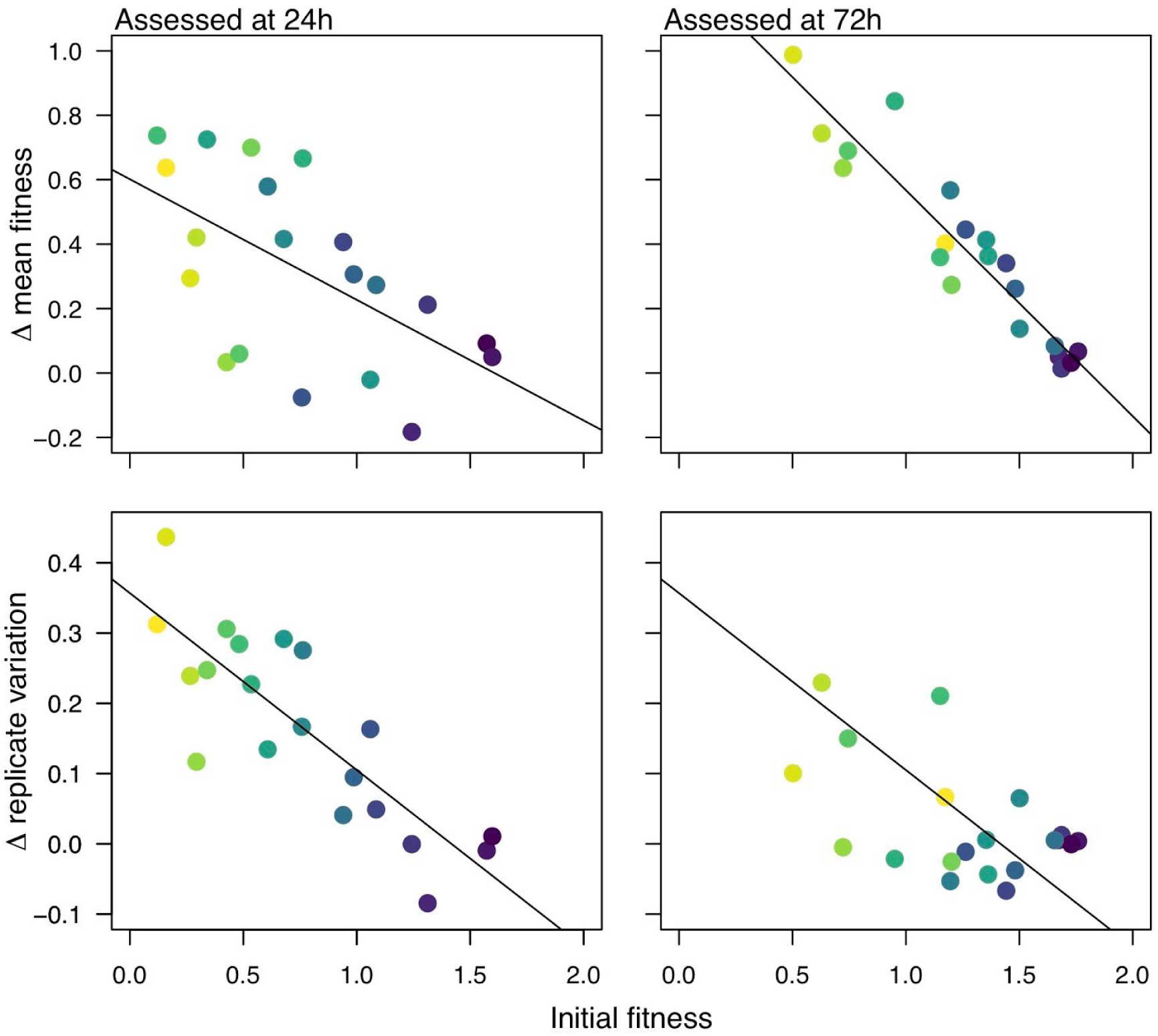
Significant association between ancestral fitness and change in mean fitness (upper graphs) and replicate variability (lower graphs) . Fitness was measured as optical density in YPD + 1 μg/mL fluconazole after 24 h (left panels) and 72 h (right panels). The colours are as in Figure 1, based on ancestral fitness at 72 h (low ancestral fitness = yellow, high ancestral fitness = purple). The regression line is for visualization purposes to illustrate the one-way relationship between ancestral fitness and change in fitness or change in variability, which was significant (p < 0.05) in multi-way models that also take into account clade and mating locus (p > 0.05 in all cases; see text for details).

### Increases in drug resistance to 1 µg/mL were common but beyond 1 µg/mL were rare

One hundred and ten replicates (76%) derived from the twelve backgrounds that had an ancestral MIC_50_ below 1 µg/mL evolved to have an MIC_50_ of 1 µg/mL within the 100 generation experiment (Figure 3a left panel). Selection for increased resistance was predominantly limited to the evolutionary drug level: only 6 replicates (4%) from four of these strains evolved a higher MIC_50_ and none of the replicates from the three strains with ancestral MIC_50_ equal to 1 µg/mL changed in MIC_50_ (Figure 3a middle) Replicates from the five strains with ancestral MIC_50_ above 1 µg/mL exhibited diverse responses (Figure 3a right panel): in three strain backgrounds (A1, A18 and A12) the replicates retained the ancestral MIC_50_; for a single strain (A20), all replicates decreased to MIC_50_ of 1 µg/mL; replicates from the fifth strain (A5) exhibited variable outcomes (four increased, four decreased, and four were unchanged in their MIC_50_). In total, only 10 replicates from five strain backgrounds increased in MIC_50_ beyond the evolutionary level of the drug (Figure 3a, Figure S1). No clear factor was associated with the appearance of resistance to drug above the evolutionary level: these events were spread across the major clades and, were not associated with mating locus (three have a homozygous mating locus, two are heterozygous). Combined, this suggests that the majority of selected mutations confer a narrow benefit up to, but not exceeding, the evolutionary level of the drug.

**Figure 3.**
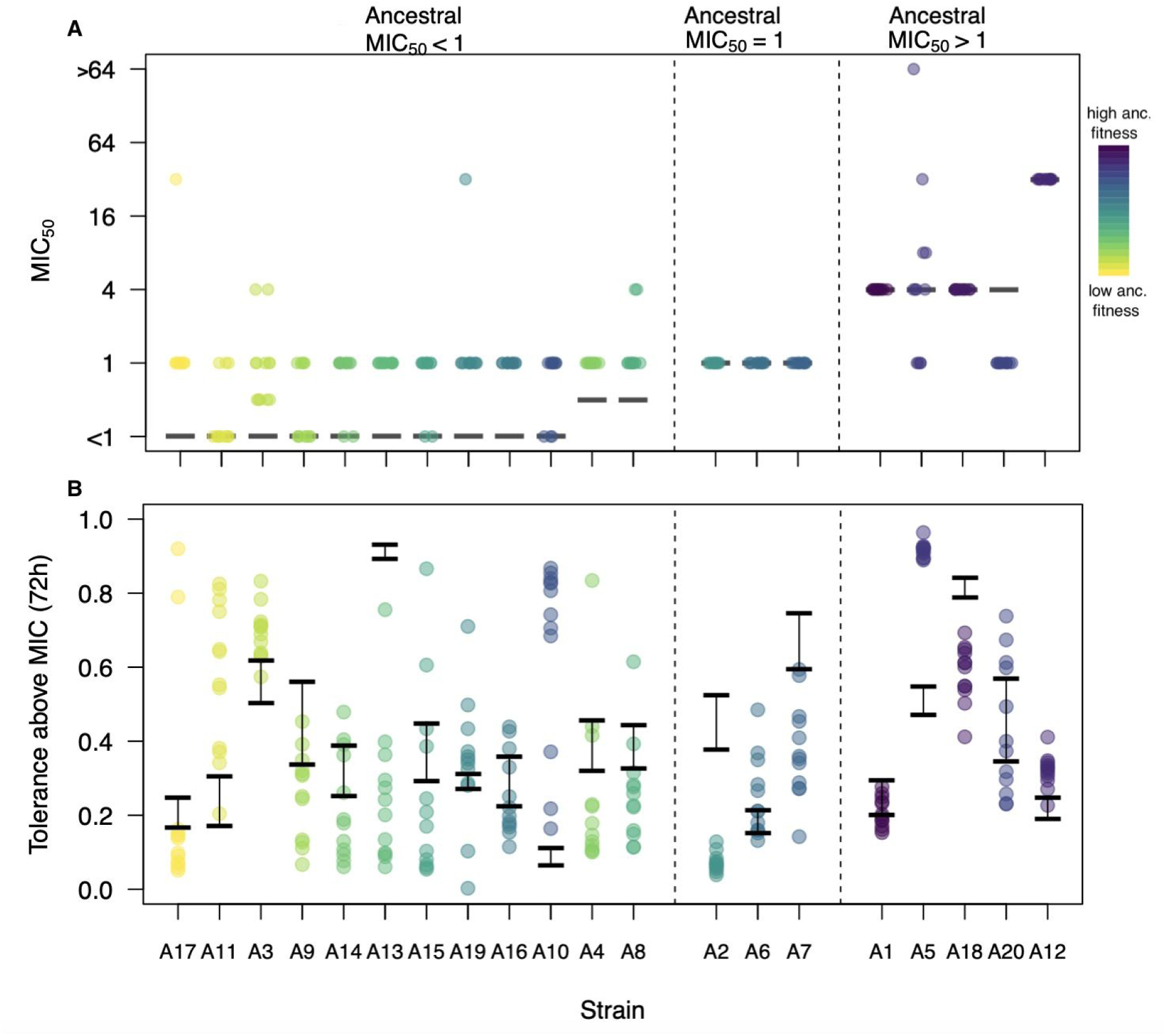
Evolved variation in clinical resistance and tolerance. Strains are arranged on the x-axis by ancestral MIC, with each panel separating the three MIC-based classes of strains. (a). The majority of evolved replicates did not acquire clinical resistance. Clinical drug resistance was measured as MIC_50_ using broth microdilution assays (1). The black line indicates ancestral MIC_50_. (b) Tolerance was variable among the replicates with evolved MIC_50_ equal to 1 (the level of drug in the evolutionary environment). The black box in panel A indicates the replicates included in panel B; two strains, A1 and A18 have no replicates with MIC_50_ = 1. Tolerance was measured as the average growth observed in supra-MIC levels of fluconazole normalized to the growth in a very low level of drug after 72 h. The black lines indicate the range of tolerance values measured among ancestral replicates. Each point represents an individually-evolved replicate line, coloured as in Figure 1, based on fitness in the evolutionary environment.

### Changes in drug tolerance were common

Changes in tolerance were prevalent among evolved replicates (Figure 3b, Figure S2). The overall picture is similar regardless of the time at which tolerance was measured (48 or 72 h, Figure S2). Consistent with a previous study examination (3), the magnitude of tolerance change increased sharply between 24 and 48 h of drug exposure; Figure S2).

Approximately 75% of the evolved replicates changed tolerance level: 55 replicates from 15 strain backgrounds increased, 122 replicates from 19 strain backgrounds decreased. The proportion of replicates within a strain that increased or decreased in tolerance varied considerably among backgrounds (Figure 3B). Ancestral strain fitness influenced the degree of stochasticity of tolerance among evolved replicates, similar to the variation in evolved fitness variation in evolved tolerance among replicates was significantly and negatively correlated with initial fitness (Spearman’s rank correlation, fitness measured at 24 h— S = 2226, p-value = 0.002, rho = -0.67; 72 h— S = 1986, p-value = 0.029, rho = -0.49). These results are consistent with the idea that genetic background, mediated in part by ancestral fitness, acts not only on evolved trait means but also on evolved trait variance.

If we consider each replicate independently, there was no association between change in tolerance and change in fitness (Linear mixed-effect model; change in fitness: F_1, 132.5_ = 0.03, *p* = 0.87, MAT zygosity: F_1,18.8_ = 0.65, *p* = 0.43, clade: F_4, 19.1_ = 1.99, *p* = 0.13). Looking at evolved replicates within each strain, a significant negative correlation between change in tolerance and change in fitness was only found for three strain backgrounds (A13, A14, and A19) (Figure S4).

### Genome size changes are pervasive

Exposure to fluconazole is known to induce the formation of tetraploid and, subsequently, aneuploid cells in *Candida albicans* (24); specific aneuploidies can provide a selective advantage under drug stress (22, 25, 48). Here, we estimated genome size by flow cytometry and found that evolved replicates underwent a significant increase in median genome size across the majority of strain backgrounds (Figure 4A, Table 2). The three exceptions were A12, the strain with which had the highest initial MIC_50_, and A4 and A9, strains with initial MIC_50_ values < 1 μg/mL. In some cases multiple subpopulations with different genome sizes were observed (see methods); for analysis, only values from the largest sub-populations were used (i.e., the most prominent peak in the flow trace), which underestimates the degree of genome size diversity in the population. Looking at A4 and A9, non-diploid sub-populations are observed for some replicates (Figure 4C, Figure S5); thus, only one strain background (A12, the strain with the highest ancestral MIC) had no replicates with a significant deviation from diploidy.

**Figure 4.**
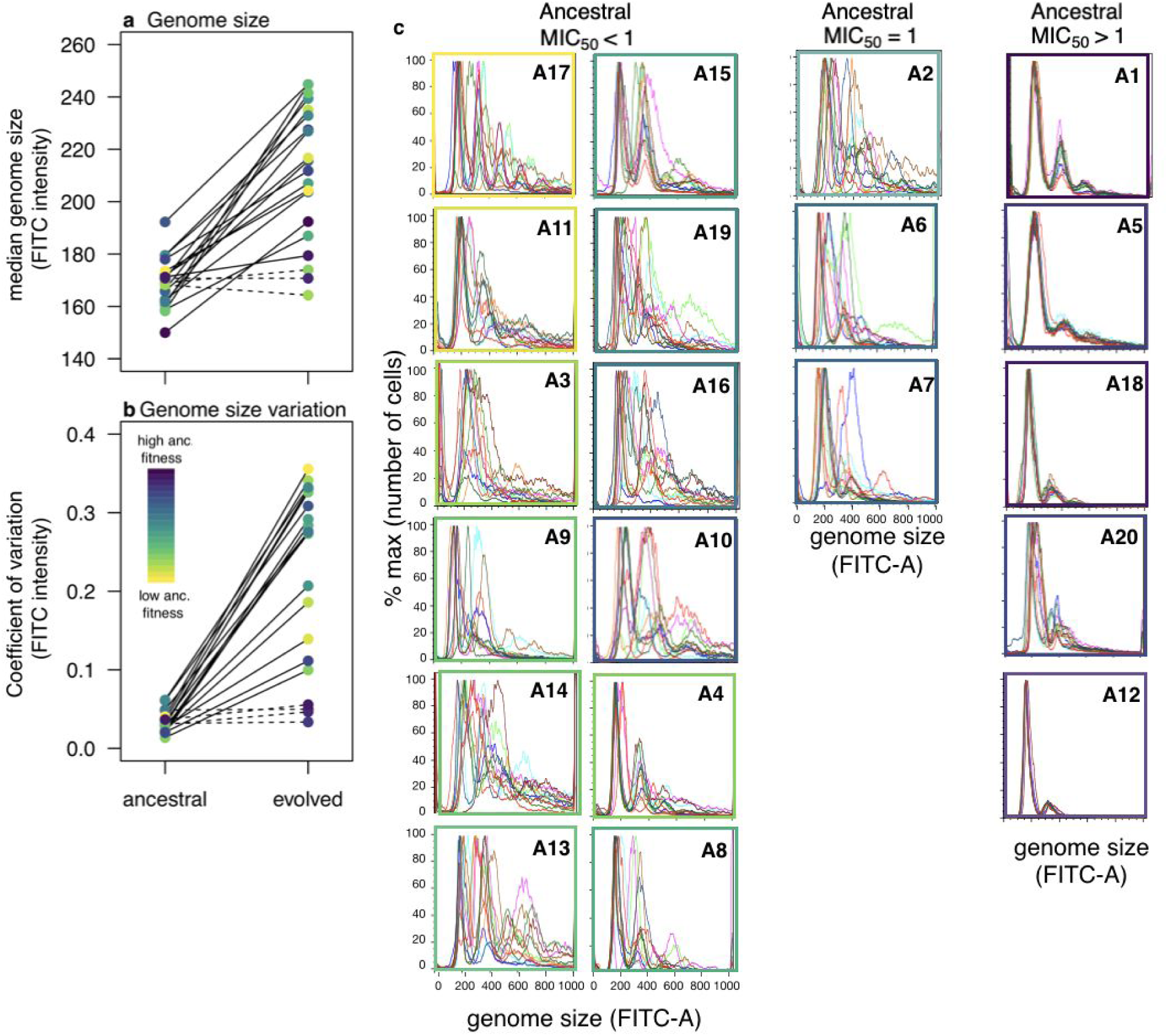
Genome size and variation in genome size increased after evolution to low fluconazole. a) Median genome size, and b) coefficient of variation (CV; i.e., variability among replicates) in each strain background. A dashed line indicates a non-significant change between ancestral and evolved replicates. c) Flow cytometry traces of each replicate evolved line, ordered and grouped by ancestral MIC50. Box colour indicates ancestral fitness in the evolutionary environment.

**Table 2.** Welch two sample t-tests to compare ancestral and evolved genome size.

| Strain | Statistics |
| --- | --- |
| A1 | $t_{20.6} = 11.70, p < 0.0001$ |
| A2 | $t_{11.2} = 5.34, p = 0.002$ |
| A3 | $t_{11.1} = 5.69, p = 0.0001$ |
| A4 | $t_{11.4} = 1.91, p = 0.08$ |
| A5 | $t_{20.14} = 18.07, p < 0.0001$ |
| A6 | $t_{11.1} = 3.82, p = 0.003$ |
| A7 | $t_{11.1} = 2.80, p = 0.017$ |
| A8 | $t_{11.1} = 2.80, p = 0.017$ |
| A9 | $t_{11.2} = 0.88, p = 0.40$ |
| A10 | $t_{11.4} = 3.31, p = 0.007$ |
| A11 | $t_{11.5} = 5.24, p = 0.0002$ |
| A12 | $t_{18.8} = 0.71, p = 0.49$ |
| A13 | $t_{11.1} = 5.46, p = 0.0002$ |
| A14 | $t_{11.1} = 4.10, p = 0.002$ |
| A15 | $t_{11.1} = 2.79, p = 0.018$ |
| A16 | $t_{11.3} = 4.17, p = 0.0015$ |
| A17 | $t_{11.1} = 2.94, p = 0.013$ |
| A18 | $t_{18.4} = 2.98, p = 0.008$ |
| A19 | $t_{11.4} = 2.93, p = 0.013$ |
| A20 | $t_{11.5} = 5.91, p < 0.0001$ |

The coefficient of variance for ploidy level, a measure of variability among replicates, also increased for the majority of strains (Figure 4B). Thus, genome size variation among replicates increased over time in drug, as expected if the drug-induced ploidy shifts and if evolved replicates acquired different final numbers of chromosomes.

Importantly, the least ploidy variation was seen with replicates evolved from the five parental strain backgrounds that had ancestral MIC_50_ levels above the evolutionary drug concentration (Figure 4C, right panels), and thus would be expected to be less sensitive to the drug stress used. When these five strains were removed from the analysis, median evolved genome size was not correlated with ancestral fitness in fluconazole (Pearson’s correlation, 24h: t_13_ = -0.30, *p* = 0.84; 72 h: t_13_ = -0.22, p = 0.83), nor with the mean fitness change in low drug (Spearman’s rank correlation, fitness at 24 h: S = 541.19, p = 0.45; 72 h: S = 953.4, p = 0.12). Genome size variation was thus extremely and equally prevalent among evolved replicates from strain backgrounds with ancestral MIC at or below the evolutionary level of drug.

These median numbers, however, obscure the tremendous variation in genome size observed among evolved replicates (Figure S5). Looking at replicates from the 15 strains with ancestral MIC_50_ ≤ 1, final genome size was not significantly correlated with any of the predictor variables that we tested (change in fitness: F_1, 46.4_ = 0.005, *p* = 0.95, clade: F_4, 9.2_ = 0.94, *p* = 0.48, MAT zygosity: F_1, 8.7_ = 0.002, *p* = 0.97). There was similarly no correlation between evolved genome size and change in fitness within replicates from any strain background (Figure S6). A14 was the only strain background with a significant negative correlation between change in tolerance and evolved genome size (Figure S7).

The variance in evolved genome size followed a similar pattern as variance in evolved tolerance: a significant negative correlation with ancestral fitness at 24 h but not at 72 h (Spearman rank correlation; 24 h fitness—S = 2076, *p* = 0.01, rho = -0.56; 72 h fitness—S = 1908, *p* = 0.05692, rho = -0.43) and a positive correlation with the variance in fitness improvement (24 h: S = 362, *p* = 0.0004, rho = 0.73; 72 h: S = 440, *p* = 0.002, rho = 0.67). Accordingly, variance in evolved genome size was also significantly correlated with variance in evolved tolerance (S = 538, p-value = 0.007, rho = 0.60). There was no direct link between evolved genome size and either fitness improvement or change in tolerance among replicates or strain backgrounds. However, the variability among evolved replicates is consistently larger in some strain backgrounds than others, regardless of the trait we measured (evolved fitness, drug tolerance, genome size), and this was mediated, in part, by ancestral strain fitness in the evolutionary environment.

## Discussion

The adaptation to antimicrobial drugs by microbial pathogens is inherently an evolutionary process that relies upon beneficial genetic mutations. To examine the influence of genetic background on adaptation in drug, we evolved 20 diverse *C. albicans* isolates (240 replicates in total) for 100 generations in 1 µg/mL fluconazole. The majority of replicates rapidly increased in fitness in the evolutionary environment, in a manner that correlated inversely with ancestral fitness. Accordingly, the majority of replicates from strains with ancestral drug susceptibility (measured as MIC_50_) below 1 µg/mL fluconazole increased MIC_50_ to the evolutionary drug level of 1µg/ml. Only ten replicates from five backgrounds increased in drug resistance beyond the evolutionary drug level, however. By contrast, changes in drug tolerance were much more common with replicates both increasing and decreasing in tolerance. These changes in tolerance may be due to pleiotropic effects from genetic changes that improve growth in the evolutionary environment, or by virtue of physiological, or possible epigenetic changes. Overall, there was little or no correlation between change in tolerance and ancestral fitness or change in fitness. However, ancestral fitness was negatively correlated with the variance in tolerance among evolved replicates, and the variance in evolved fitness was also negatively correlated with the variance in evolved genome size.

The negative correlation between ancestral fitness and the improvement in fitness, and between ancestral fitness and the variation among replicates in fitness, drug tolerance, and genome size are both consistent with predictions from Fisher’s abstract geometric model (49, 50). These results among very diverse ancestral strains are similar to the pattern of diminishing-returns for fitness seen in bacteriophage (51) and *E. coli* (52) and are consistent with the negative correlation between ancestral fitness and the rate of adaptation in strains that differ by many mutations (53). and in strains differ by only one or a few beneficial (54, 55) or deleterious mutations (56).

Only a small number of replicates increased MIC_50_ to a drug level above the evolutionary environment. From the raw (optical density) data for specific replicates from A3, A8, A17 and A19 (ancestral MIC_50_ of all four < 1 µg/mL), MIC_50_ clearly increased beyond 1 µg/mL (Figure S1), indicating that these isolates acquire resistance to a higher drug concentration. Conversely, replicates from strain A20 (ancestral MIC_50_ = 4 µg/mL) uniformly improved their fitness at 1 µg/mL fluconazole, yet had reduced fitness at 4 µg/mL fluconazole, indicative of a cost-benefit trade-off for fitness in lower vs. higher drug concentrations of fluconazole. Other MIC_50_ results require a more cautioned biological interpretation and may be due to a technical issue, rather than biological improvement. MIC_50_ is calculated as drug concentration where growth is reduced 50% relative to growth in no drug; hence an alternative mechanism to (numerically) increase the measured MIC_50_ is to reduce growth in the absence of drug. This appears to have occurred for some replicates, primarily from initially low-MIC_50_ ancestral strains (i.e., single replicates in A3 and A8, Figure S1). Replicates from strain A5 highlight how small differences in OD can influence the calculation of MIC_50_: four strains have numerically increased and four have numerically decreased MIC_50_, yet the actual numbers reveal only a minor separation among replicates (Figure S1). Visual examination of both the assay plates and graphic display of the raw optical density values is thus required to properly interpret numerical differences in MIC_50_

In bacteria, exposure to subinhibitory concentrations of antibiotics can select for *de novo* mutations that confer resistance (57–62) or provide a pleiotropic benefit alongside existing resistance mutations (e.g., 57, 63–66). Here, consistent patterns among strains are lacking: of the five strains backgrounds that were exposed to a subinhibitory level of fluconazole, no significant changes were observed among replicates from three of them (A1, A12, A18), MIC_50_ decreased in all replicates from one replicate (A20), and variable, yet numerically significant, changes occured in replicates of another strain (A5). This variability in phenotypic outcomes is reminiscent of the case in *Sclerotinia sclerotiorum,* a plant pathogenic fungus, where both MIC increases and decreases (and no change) were observed and no consistent relationship was found between the change in resistance and exposure to sublethal concentrations of five different antifungal drugs (67). Fungal strains exposed to subinhibitory drug levels thus seem less likely than bacteria to gain an advantage at higher drug levels. Given the possible role of aneuploidy in this process, we speculate that differences in chromosome genometry (circular vs. linear) and mechanisms that affect chromosome segregation may underlie these differences in the dynamics of antimicrobial responses.

The majority of replicates from strains with ancestral MIC at or below the evolutionary drug concentration had increased DNA content, interpreted as larger genome size. Furthermore, changes in DNA content and changes in fitness, MIC_50_, or tolerance did not correlate, consistent with the idea that if the fluconazole-exposed isolates carry several aneuploid chromosomes, not all of them are necessarily causative of the observed fitness increases. Exposure to fluconazole at 10 µg/ml is known to induce the formation of ‘trimeras’ (24), cells indicative of mitotic defects that result in aneuploidy at higher fluconazole concentrations; trimeras also were evident when lab strain SC5324 was exposed to 1 µg/ml fluconazole (Maayan Bibi and Judith Berman, unpublished data). Growth in fluconazole is expected to exert selection pressure for some aneuploids more than others; specific genes within an aneuploid chromosome that are responsible for increased drug resistance (26) and drug tolerance (27) also have been identified. In many other cases, more than one gene may contribute to the phenotype (22, 27, 48, 68, 69); (22, 27, 48, 68). While we assume that changes in DNA content of a given isolate are largely due to chromosomal aneuploidy, it is important to note that increased levels of mitochondrial and/or ribosomal DNA can also contribute to differences in DNA content level detected by flow cytometry, as was recently shown to occur in different deletion mutants within the ‘isogenic’ collection of *S. cerevisiae* deletion mutants (70).

Given that some gene products, and the allelic ratios of the genes encoding them, may be more limiting than others in the face of a specific stress, it is quite clear that the degree to which a given aneuploid chromosome (or other copy number change) may contribute to stress tolerance or resistance is likely to be affected by the genetic background. The lack of a correlation between evolved genome size and fitness also suggests that there is a low cost to aneuploidy in this environment (i.e., there is no clear benefit to increased size, but there is also no clear cost). We observed lower genome size variation in some strain backgrounds than others of a similar ancestral fitness (i.e., A4, A9), suggesting that these strain backgrounds may be less tolerant of aneuploidy than others (15). Variability in DNA content among evolved replicates was also very low for strains with ancestral MIC_50_ > 1 µg/mL, presumably because these cells were under little stress in the evolutionary environment (Figure S5). Thus, we posit that the observed variation in evolved ploidy may be integrally connected to the rapid appearance of altered drug responses. This observation may have clinical relevance, given that aneuploidy was common, albeit transient, in a study of sequential clinical isolates of *C. albicans*. Importantly, aneuploidy appeared concomitant with major shifts in drug resistance, yet was not retained in strains that acquired *bona fide* drug resistance (13). The DNA content measurements were captured directly from populations of cells in the drug environment, and it is important to note that aneuploid chromosomes can be lost extremely rapidly in the absence of selective pressure (e.g., following a single overnight growth cycle in permissive medium). This highlights the important idea that aneuploidy may provide a rapid, highly frequent, yet transient and suboptimal, genome change that facilitates adaptation until more robust and stable genome changes (e.g., point mutations) can be acquired (71, 72).

## Conclusion

Experimental evolution studies can isolate important factors that influence adaptation. Here, genetic background had a significant influence on the rate and variability of adaptation, mediated in part through ancestral fitness relative to the selective conditions used. Evolved changes in DNA content were prevalent among strains with ancestral MIC < 1, and largely absent from those with ancestral MIC above the condition used for evolution, highlighting the context-dependence (relative to strain MIC) of drug stress. Importantly, ancestral fitness was correlated with evolved variation among replicates for fitness, drug tolerance, and genome size, thereby emphasizing that strain background can influence both the magnitude and variation in adaptive responses to drug.

## Materials and Methods

### Strains

Twenty clinical strains of *Candida albicans* were selected to represent the phylogenetic diversity of the species. The strain set includes at least four strains from each of the four major clades that encapsulate ∼70% of the typed *C*. *albicans* strains and spans nearly the entire known phylogenetic diversity of the species (47, 73) as well as four commonly-studied laboratory strains (SC5314, FH1, DSY294, T188) (Table 1). For each clade, strains with both heterozygous (*MAT**a**/MATα*) and homozygous (*MAT**a/**MAT**a***, *MATα/MATα*) mating loci were chosen. All 20 strains were ancestrally diploid though strain A1 was trisomic for chr7, and A11 was trisomic for chr4. Strains were chosen blind with resplect to ancestral fitness or drug resistance. Full strain information including clade designation, country of origin, and infection niche were obtained from the original manuscripts (Table 1) and (74). Mating type genotype was confirmed by PCR with *MAT**a*** and *MATα* specific primers (*MAT**a*** F-TTGAAGCGTGAGAGGCAGGAG, *MAT**a*** R-GTTTGGGTTCCTTCTTTCTCATTC, *MATα* F-TTCGAGTACATTCTGGTCGCG, *MATα* R-TGTAAACATCCTCAATTGTACCCG). All strains were initially streaked onto YPD and grown for 48h at 30°C. A single colony was frozen down in 15% glycerol and stored at –80 °C. Thus, minimal genetic variation should be present in the initial freezer stock, which we refer to throughout as the “ancestral strains”.

### Evolution Experiment

Strains were evolved in 1 μg/mL fluconazole, the epidemiological cut-off value that denotes the upper limit of drug susceptibility (MIC_50_) in the wild type *C. albicans* population (5). This concentration of drug was equivalent to the ancestral MIC_50_ of three strains, above the MIC_50_ for 12 strains and below the MIC_50_ for one highly resistant strain (MIC_50_ = 32) and four additional strains (MIC_50_ = 4) (Table 1).

To initiate the evolution experiment we generated twelve independent replicates from each ancestral strain. Cultures were struck from frozen ancestral stocks onto YPD plates (1% yeast extract, 1% peptone, 2% dextrose, 1% agar; the standard lab rich medium) and incubated at 30°C overnight. For each strain, colonies were randomly chosen by spotting each plate twelve times and picking the closest colony to each dot. Colonies were separately inoculated into 1 mL YPD in a 96-well (3 mL) deep culture box and grown shaking overnight at 30 °C.

From the overnight cultures, we froze 100 μL from each replicate in duplicate in 50% glycerol as the ancestral replicates. Overnight cultures were diluted 1:1000 into YPD+fluconazole in two steps: first, 10 μL of the overnight culture was transferred into 990 μL YPD (a 1:100 dilution), followed by a transfer of 20 uL diluted culture into 180 μL of YPD + 1.11 μg/mL FLC in round-bottom microtiter plates. To minimize the likelihood of contamination and keep environmental conditions similar, culture from replicates from one strain was inoculated into row A while culture from replicates from a second strain was inoculated into row H. Plates were sealed with breathe-easy sealing membranes (Sigma Z380058) and incubated static at 30 °C to mimic the static growth used for clinical resistance assays. Plates were contained within small sealed Rubbermaid containers with wet paper towels inside to minimize evaporation.

After 72 hours, wells were mixed by pipetting and another two-step 1:1000 transfer was conducted into fresh YPD + fluconazole medium. In total 10 transfers were conducted, yielding 100 generations of evolution (9.97 generations between transfers, log_2_(1000) = 9.97 * 10 transfers = 99.7 generations). 50µl of the evolved replicate cultures were frozen in duplicate in 50% glycerol after the tenth transfer and maintained at -80 °C.

### Growth in the Evolutionary Environment

We measured fitness in the evolutionary drug environment as optical density (A_600_) at both 24 and 72 h. Optical density reflects the ability to convert nutrients from the environment into cellular biomass. Fitness at 24 h can be thought of as a composite parameter reflecting both lag phase and the exponential growth rate (strains that have either a lower lag or a faster growth rate will have higher 24 h fitness; (75). Fitness at 72 h reflects growth at the time of transfer and the amount of biomass present at stationary phase. OD at 24 h is also consistent with the clinical assessment of drug resistance.

### Clinical Resistance & Tolerance

The initial susceptibility and tolerance of all strains were tested using broth microdilution liquid assays to measure the minimal inhibitory concentration (MIC_50_) and tolerance as supraMIC growth (SMG), respectively. The liquid assay experiments followed the initial cell dilution regulations from the clinical CLSI M27-A guidelines (http://clsi.org/), except with YPD incubated at 30 °C as the base medium and optical density at A_600_ (OD) instead of a McFarland 0.5 standard to determine the initial density of cells. OD readings were taken at 24 hours after inoculation to calculate MIC_50_ (the point at which optical density was reduced by 50% compared to optical density in YPD without drug).

Four liquid broth microdilution assays were conducted on both the ancestral and evolved replicates, and an additional two assays were conducted on the ancestral replicates. We were not able to assay all drug concentrations in each assay due to capacity (2 time points x 20 strain backgrounds x 12 replicates). A single measurement was taken for each replicate at each concentration of drug measured in a given experiment (see Table S2). The median OD among experiments was determined at each concentration of drug for each replicate. Following guidelines, the MIC_50_ was then calculated as the highest concentration of drug with an OD greater than 50% of the measured OD in YPD (i.e., optical density in medium without drug).

Tolerance was measured from the liquid assay results as in (3). The average growth (measured as optical density, A_600_) in the concentrations of drug that were measured above the MIC was determined for each ancestral and evolved replicate. This was divided by the measured growth in the lowest drug level (0.0625 uM fluconazole) so that tolerance reflects the fraction of realized growth and is between 0 and 1. Tolerance was assessed at 24, 48 and 72 h. We consider evolved replicates to have increased or decreased in tolerance when their measured tolerance was respectively above or below the range of ancestral tolerance levels for replicates from that strain.

### Ploidy variation

Flow cytometry was performed on a BD Biosciences BD LSR II. In all cases, all replicates from the same time point were fixed, stained and measured in parallel. Ancestral replicates were measured twice independently from the freezer stocks maintained at -80 °C. Freezer cultures were thawed, mixed, and 10 μL was added to 500 μL, YPD in deep 96-well boxes, covered with a breathe-easy membrane, and shaken at 200 RPM, 30 °C overnight. After ∼ 16 hours of growth, 20 μL of culture was washed in 180 μL of TE, in a round bottom microtiter plate, pelleted, and resuspended in 20 μL TE and fixed by adding 180 μL 95% cold ethanol.

Samples from evolved replicates were fixed in ethanol at the end of the last growth cycle; 50 μL of 72 h culture from t10 was washed in 150 μL TE in a round bottom microtiter plate, pelleted and resuspended in 20 μL TE and 180 μL 95% cold ethanol. Ethanol-fixed cultures were stored at -20 °C for up to 4 weeks. The remainder of the protocol was identical for both time points, following Gerstein et al. 2017. As described in detail in Gerstein *et al.* ((76)) we used the cell cycle analysis in Flow-Jo (Treestar) to determine the mean G1 peak for each replicate; when more than one peak was evident we recorded both the major and minor G1 peaks.

Although we always used the same machine settings, subtle but significant variation is always observed in flow cytometry data. To better compare between t0 and t10 data we performed a day-correction based on the median G1 intensity of the A12 ancestral and evolved replicates, which always measured cleanly as diploids.

### Statistical methods

All analysis was conducted in the R Programming Language (77). To maximize statistical power when testing the influence of mating type, we examined the effect of a heterozygous mating type vs. homozygous mating type, i.e., we combined *MATa/a* and *MATα/α* strains.

MIC_50_ values were log transformed prior to statistical analysis. When parametric tests were used, all assumptions were tested and met. When data transformations were insufficient to meet the test assumptions, nonparametric tests were used. Spearman’s rank correlation was used when comparing the mean responses among replicates. This correlation method uses a rank-based measure which does not require the replicate data to come from a bivariate normal distribution. In all cases the specific test is indicated inline.

We used linear-mixed effect models to determine which factors influenced evolved tolerance and genome size. Since there was significant variation in ancestral tolerance, the response variable in the tolerance model was evolved - ancestral tolerance. In both models the predictor variables were change in fitness (OD at 72 h, evolved - ancestral), zygosity and clade, with strain as a random effect. The models were implemented with the lmer package (78) in the R programming language (77). Significance was determined from the Type III ANOVA Table with the Satterthwaite’s approximation to degrees of freedom.

## Acknowledgements and Funding Information

We thank R Urbitas, D Abbey, MA Hickman, M McClellan, E Shtifman Segal, D Tank and the University of Minnesota Flow Cytometry core facility for technical support, A Selmecki for thoughtful discussions and J Hill for helpful comments on an early version of the manuscript. This work was supported by a European Research Council (Advanced Award 340087, RAPLODAPT to J.B.). ACG is grateful to the Azrieli Foundation for the award of an Azrieli Postdoctoral Fellowship, and for support from a National Sciences and Engineering Research Council of Canada Postdoctoral Fellowship and a Banting Fellowship from the Canadian Institutes of Health Research.

## Author Contributions

All authors conceptualized the study and wrote the manuscript. ACG conducted the research and analyses.

## Data Accessibility

All data and the R code required to run the analyses and create the visualizations are available on GitHub (https://github.com/acgerstein/C_albicans-LDE).

## Supplementary Tables and Figures

**Table S1.** The majority of strains evolved an increased growth ability in the evolutionary environment. The test column indicates the test that was run: a t-test when both ancestral and evolved replicate groups were normally-distributed with equal variance, a t-test with welch approximation for degrees of freedom when variances were unequal, or the Wilcoxon Rank Sum test when the data from at least one group was not normally distributed. Equal variance was assessed with an F test, normality with the Shapiro-Wilk test (assumption test results not shown).

| Line | Growth ability at 24 h |  |  |  |  | Growth ability at 72 h |  |  |  |  |
| --- | --- | --- | --- | --- | --- | --- | --- | --- | --- | --- |
|  | test | t-value | df | p | evol-anc | test | t-value | df | p | evol-anc |
| A1 | t-test | -6.85 | 22 | <0.0001 | 0.09 | t-test | -12.49 | 22 | <0.0001 | 0.07 |
| A2 | wilcox | 89 | - | 0.35 | -0.02 | wilcox | 0 | - | <0.0001 | 0.45 |
| A3 | wilcox | 0 | - | <0.0001 | 0.42 | t-test | -12.96 | 22 | <0.0001 | 0.64 |
| A4 | welch | -9.02 | 11.47 | <0.0001 | 0.7 | wilcox | 0 | - | <0.0001 | 0.84 |
| A5 | welch | -5.58 | 13.31 | 0.0001 | 0.21 | t-test | -8.49 | 22 | <0.0001 | 0.05 |
| A6 | t-test | -4.29 | 22 | 0.0003 | 0.27 | wilcox | 0 | - | <0.0001 | 0.34 |
| A7 | t-test | -4.17 | 22 | 0.0004 | 0.31 | t-test | -7.95 | 22 | <0.0001 | 0.26 |
| A8 | welch | -6.67 | 11.79 | <0.0001 | 0.67 | welch | -19.22 | 15.01 | <0.0001 | 0.57 |
| A9 | wilcox | 96 | - | 0.18 | 0.03 | welch | -11.17 | 12.63 | <0.0001 | 0.69 |
| A10 | welch | 1.03 | 13.14 | 0.32 | -0.08 | wilcox | 7 | - | <0.0001 | 0.14 |
| A11 | welch | -3.58 | 11.5 | 0.0041 | 0.29 | welch | -8.09 | 12.45 | <0.0001 | 0.74 |
| A12 | t-test | 7.26 | 22 | <0.0001 | -0.18 | wilcox | 36 | - | 0.0387 | 0.01 |
| A13 | welch | -7.45 | 11.16 | <0.0001 | 0.74 | welch | -10.56 | 15.23 | <0.0001 | 0.4 |
| A14 | wilcox | 94 | - | 0.22 | 0.06 | wilcox | 21 | - | 0.0023 | 0.36 |
| A15 | welch | -7.35 | 12.41 | <0.0001 | 0.72 | t-test | -9.91 | 22 | <0.0001 | 0.27 |
| A16 | welch | -8 | 14.74 | <0.0001 | 0.58 | welch | -21.39 | 12.27 | <0.0001 | 0.36 |
| A17 | welch | -4.77 | 11.07 | 0.0006 | 0.64 | welch | -18.68 | 14.55 | <0.0001 | 0.99 |
| A18 | t-test | -4.92 | 22 | 0.0001 | 0.05 | t-test | -6.34 | 22 | <0.0001 | 0.03 |
| A19 | welch | -3.9 | 11.83 | 0.0022 | 0.42 | t-test | -12.61 | 22 | <0.0001 | 0.41 |
| A20 | t-test | -9.59 | 22 | <0.0001 | 0.41 | t-test | -11.5 | 22 | <0.0001 | 0.08 |

**Table S2.** Minimum inhibitory concentration experiments.

| Exper | Time | Fluconazole level (μg) |  |  |  |  |  |  |  |
| --- | --- | --- | --- | --- | --- | --- | --- | --- | --- |
|  |  | 0 | 0.5 | 1 | 4 | 8 | 32 | 128 | 512 |
| 1 | Anc&Evol | x |  | x |  | x | x |  |  |
| 2 | Anc | x |  | x |  |  |  |  |  |
| 3 | Anc&Evol | x | x | x | x |  |  |  |  |
| 4 | Anc | x |  | x | x | x | x |  |  |
| 5 | Anc&Evol | x |  |  |  | x | x | x |  |
| 6 | Anc&Evol | x |  |  |  |  | x | x | x |

**Figure S1.**
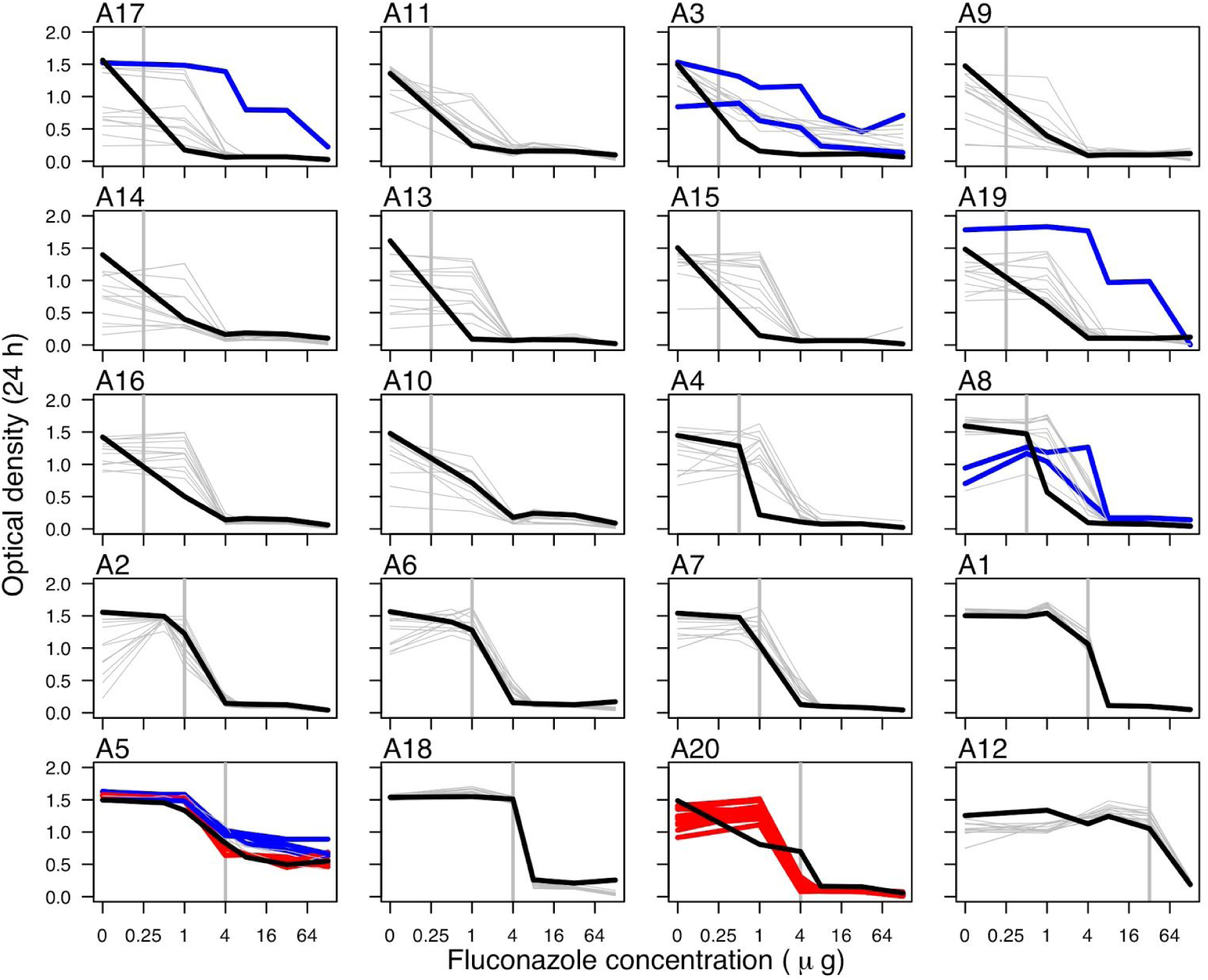
Raw broth microdilution (MIC) data at 24 h. Strains are arranged based on ancestral MIC_50_ (calculated as the drug concentration at which a 50% reduction of growth is observed after 24 h compared to growth in no drug), indicated by the horizontal grey line in each panel. The thick black line indicates the mean ancestral trace from 12 replicates. The other lines indicate evolved replicates that that increased MIC_50_ (thick blue lines), decreased MIC_50_ (thick red lines) or did not change (grey lines).

**Figure S2.**
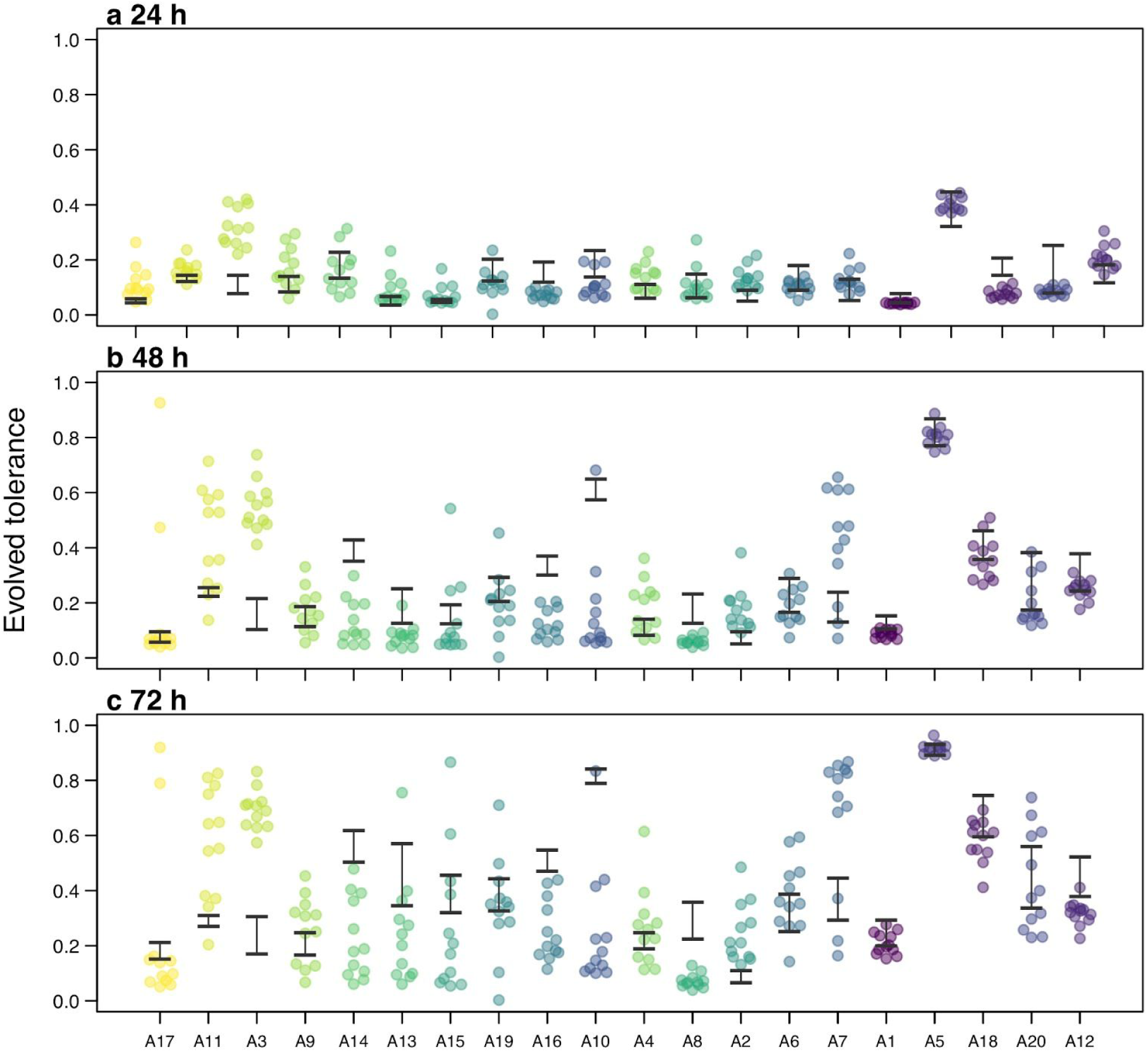
Tolerance of evolved strains measured at a) 24 h, b) 48 h, and c) 72 h. Tolerance was measured as the growth observed in supra-MIC levels of fluconazole (as appropriate to each replicate) normalized to the growth in the absence of drug. Strains are arranged on the x-axis (and coloured) by initial growth in the evolutionary environment. Each point represents an individually-evolved replicate line. The black lines indicate the range of tolerance values measured among ancestral replicates.

**Figure S3.**
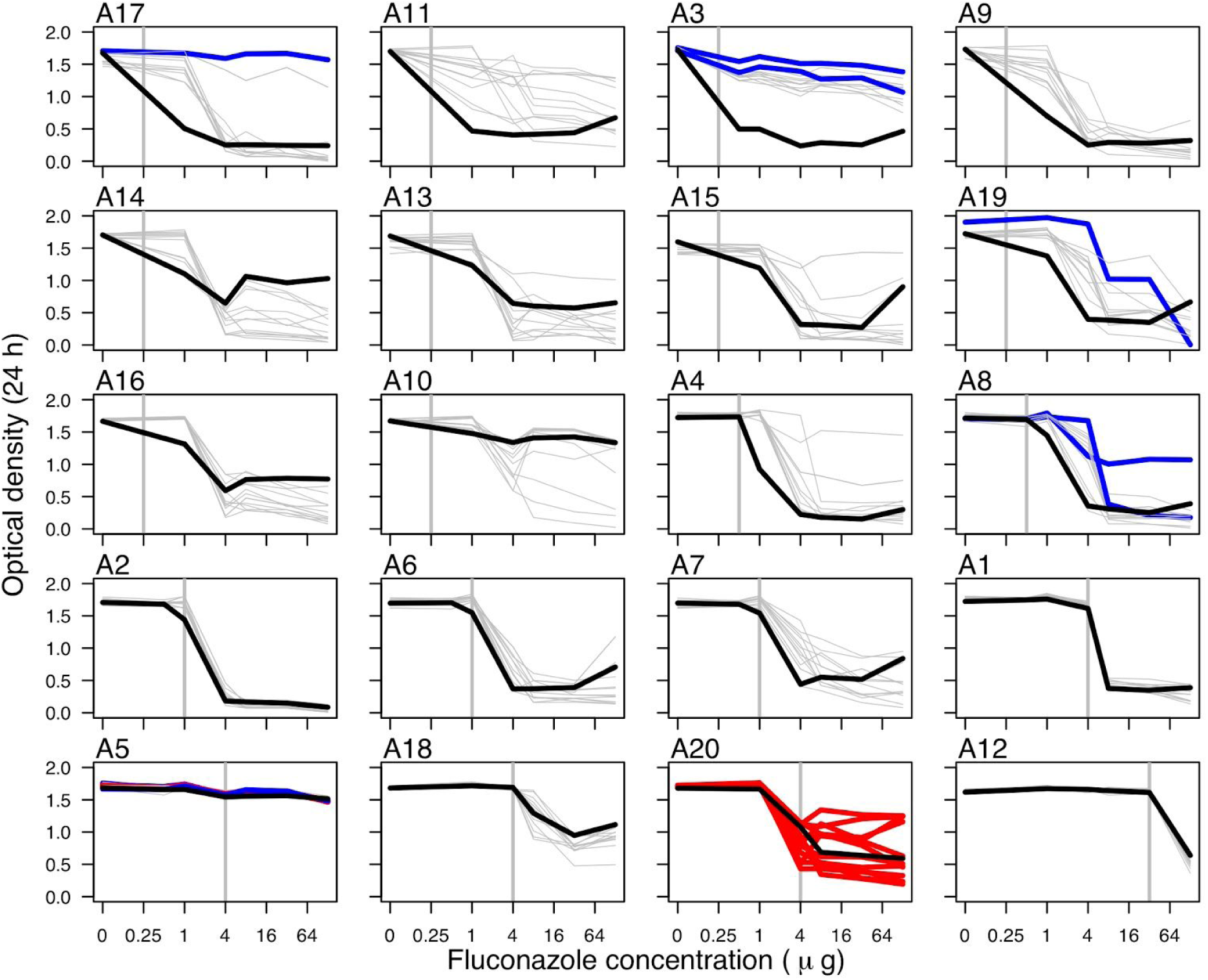
Raw broth microdilution data at 72 h. Strains are arranged based on ancestral MIC_50_ (calculated as the drug concentration at which a 50% reduction of growth is observed after 24 h compared to growth in no drug), indicated by the horizontal grey line in each panel. The thick black line indicates the mean ancestral trace of optical density measurement at 72 h from 12 replicates. The other lines indicate evolved replicates that that increased MIC_50_ (thick blue lines), decreased MIC_50_ (thick red lines) or did not change (grey lines).

**Figure S4.**
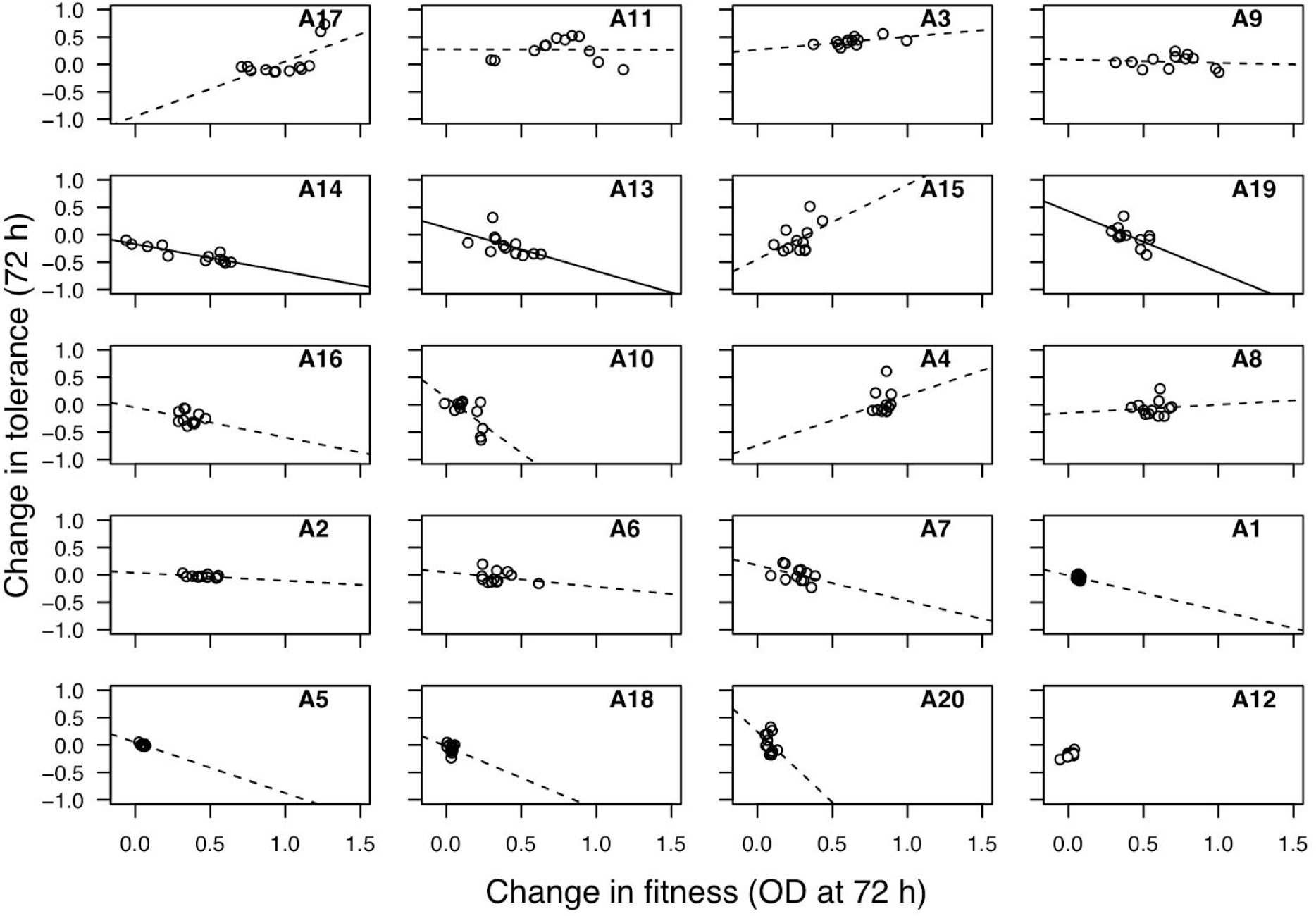
Evolved fitness and evolved tolerance are not consistently correlated among replicates. Fitness was measured as optical density after 72 hours in low fluconazole. Tolerance was measured as the average growth observed in supra-MIC levels of fluconazole normalized to the growth in a very low level of drug after 72 h (“SMG”). The black line indicates the correlation among all replicates; a solid line indicates a significant correlation (*p* < 0.05).

**Figure S5.**
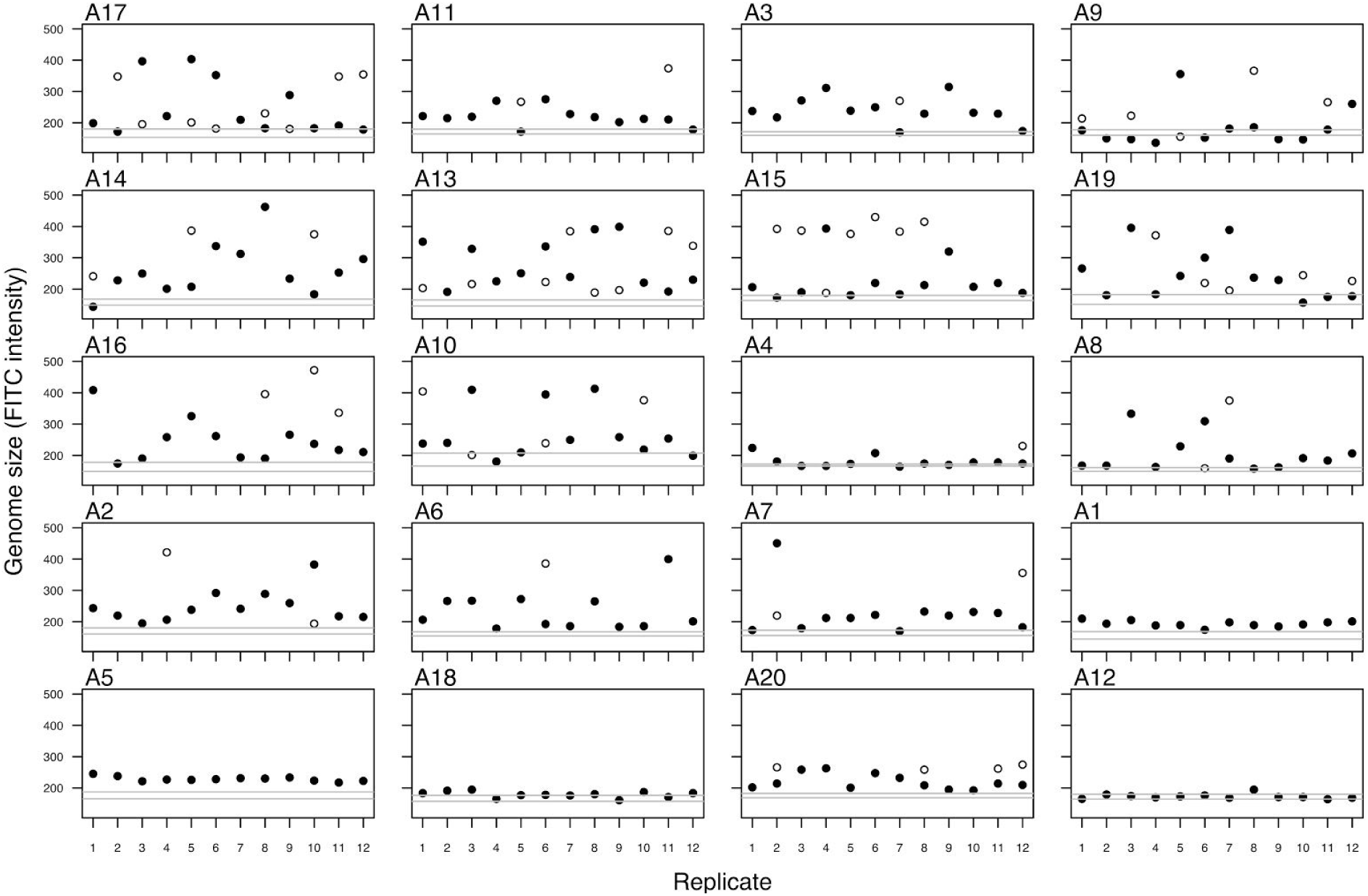
The majority of evolved replicates increased genome size. Filled circles indicate the most prominent evolved genome size peak; when multiple G1 peaks were present this is indicated with an open circle. Strains are ordered based on ancestral MIC. The grey lines in each panel indicate the range of genome size values measured for the 12 ancestral replicates from each strain at the first transfer.

**Figure S6.**
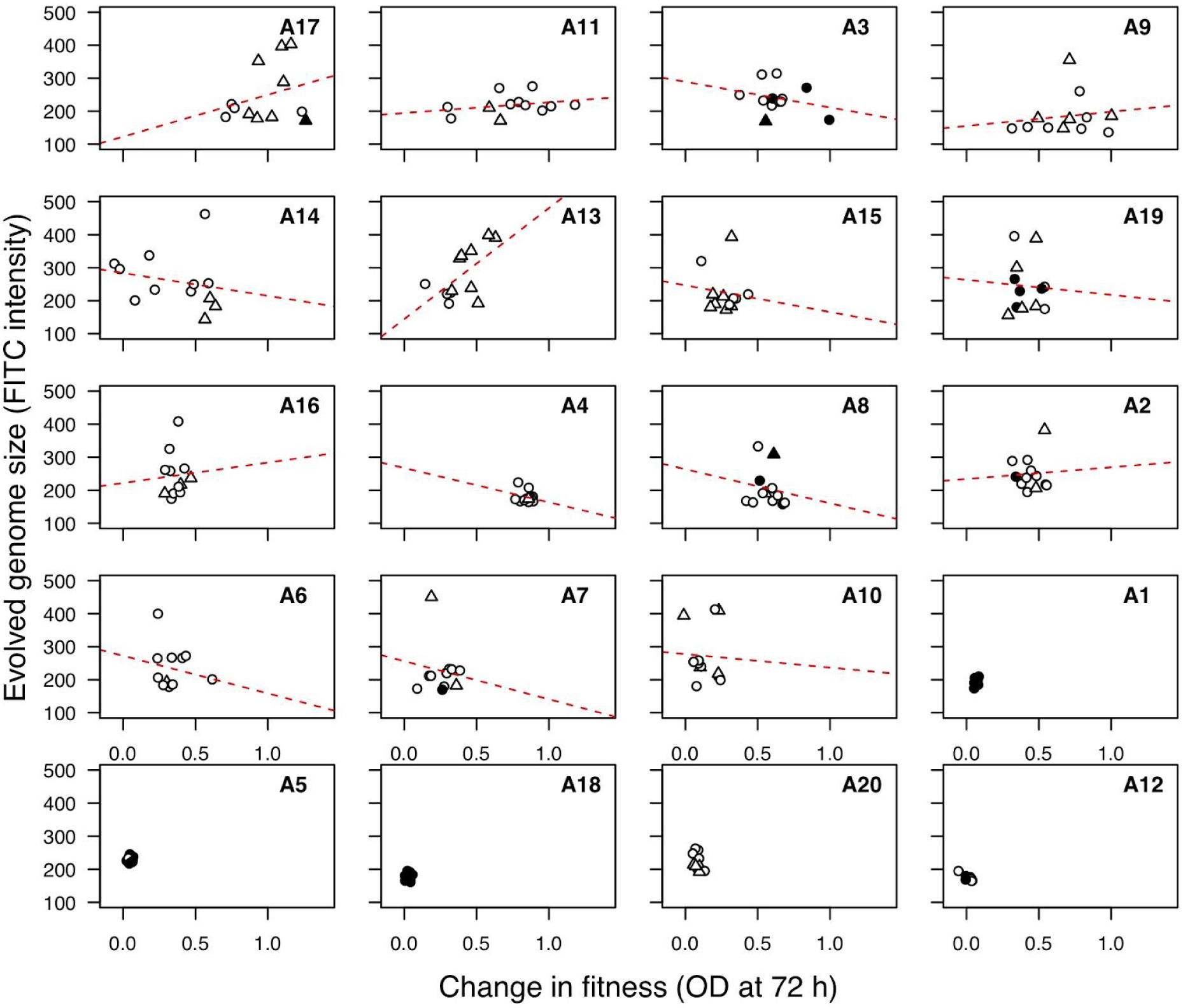
Evolved genome size and change in fitness are not correlated. Evolved genome size indicates the most prominent G1 peak; when multiple G1 peaks were present this is indicated with a triangle symbol. Fitness was measured as optical density after 72 hours in fluconazole. Each point represents an independently evolved replicate line. Filled-points are those lines that have an MIC_50_ > 1. Strains are ordered based on ancestral fitness. A correlation was not assessed for the five strains that had ancestrally high MIC levels above the evolutionary environment and did not evolve variation in genome size.

**Figure S7.**
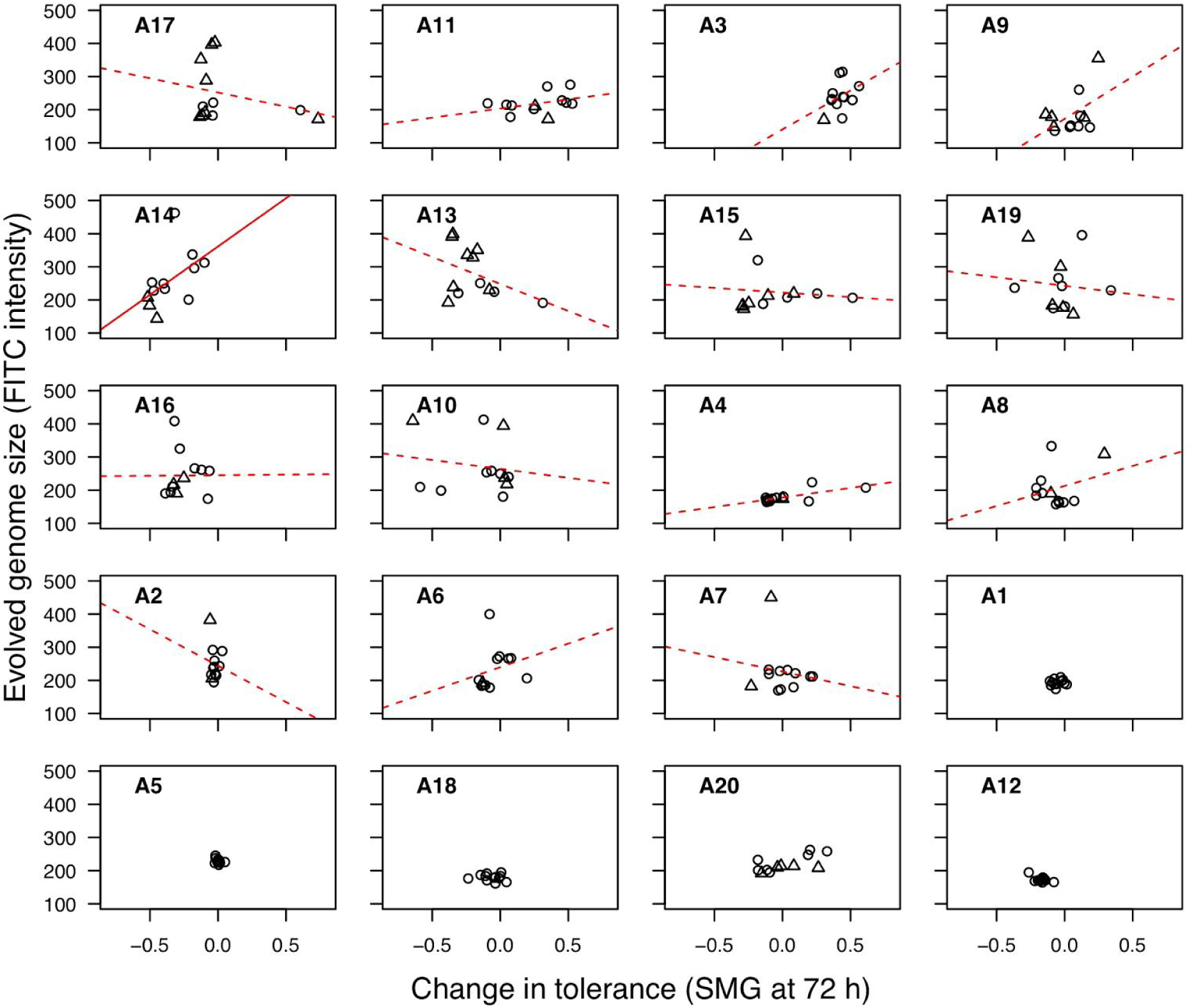
Evolved genome size and change in tolerance are not generally correlated. Evolved genome size indicates the most prominent G1 peak; when multiple G1 peaks were present this is indicated with a triangle symbol. Tolerance was measured as the average growth observed in supra-MIC levels of fluconazole normalized to the growth in a very low level of drug after 72 h (“SMG”). Each point represents an independently evolved replicate line. Strains are ordered based on ancestral fitness. A solid line indicates a significant (p < 0.05) spearman’s correlation, the dashed lines are provided for visualization. A correlation was not assessed for the five strains that had ancestrally high MIC levels above the evolutionary environment and did not evolve variation in genome size.

